# Origin and Evolution of a Pandemic Lineage of the Kiwifruit Pathogen *Pseudomonas syringae* pv. *actinidiae*

**DOI:** 10.1101/085613

**Authors:** Honour C. McCann, Li Li, Yifei Liu, Dawei Li, Pan Hui, Canhong Zhong, Erik Rikkerink, Matthew Templeton, Christina Straub, Elena Colombi, Paul B. Rainey, Hongwen Huang

## Abstract

Recurring epidemics of kiwifruit (*Actinidia* spp.) bleeding canker disease are caused by *Pseudomonas syringae* pv. *actinidiae* (*Psa*), whose emergence coincided with domestication of its host. The most recent pandemic has had a deleterious effect on kiwifruit production worldwide. In order to strengthen understanding of population structure, phylogeography and evolutionary dynamics of *Psa*, we sampled 746 *Pseudomonas* isolates from cultivated and wild kiwifruit across six provinces in China, of which 87 were *Psa*. Of 234 *Pseudomonas* isolated from wild *Actinidia* spp. none were identified as *Psa*. Genome sequencing of fifty isolates and the inclusion of an additional thirty from previous studies show that China is the origin of the recently emerged pandemic lineage. However China harbours only a fraction of global *Psa* diversity, with greatest diversity found in Korea and Japan. Distinct transmission events were responsible for introduction of the pandemic lineage of *Psa* into New Zealand, Chile and Europe. Two independent transmission events occurred between China and Korea, and two Japanese isolates from 2014 cluster with New Zealand *Psa*. Despite high similarity at the level of the core genome and negligible impact of within-lineage recombination, there has been substantial gene gain and loss even within the single clade from which the global pandemic arose.

**SIGNIFICANCE STATEMENT:** Bleeding canker disease of kiwifruit caused by *Pseudomonas syringae* pv. *actinidiae* (*Psa*) has come to prominence in the last three decades. Emergence has coincided with domestication of the host plant and provides a rare opportunity to understand ecological and genetic factors affecting the evolutionary origins of *Psa*. Here, based on genomic analysis of an extensive set of strains sampled from China and augmented by isolates from a global sample, we show, contrary to earlier predictions, that China is not the native home of the pathogen, but is nonetheless the source of the recent global pandemic. Our data identify specific transmission events, substantial genetic diversity and point to non-agricultural plants in either Japan or Korea as home to the source population.

## INTRODUCTION

A pandemic of kiwifruit (*Actinidia* spp.) bleeding canker disease caused by *Pseudomonas syringae* pv. *actinidiae* (*Psa*) emerged in 2008 with severe consequences for production in Europe, Asia, New Zealand and Chile (1-7). Earlier disease epidemics in China, South Korea and Japan had a regional impact, however, as infections were often lethal and the pathogen rapidly disseminated, it was predicted to pose a major threat to global kiwifruit production (8, 9). Despite recognition of this threat - one subsequently realized in 2008 - little was done to advance understanding of population structure, particularly across regions of eastern Asia that mark the native home of the genus *Actinidia*.

The origins of agricultural diseases and their link with plant domestication is shrouded by time, as most plant domestication events occurred millennia ago. Kiwifruit (*Actinidia* spp.) is a rare exception because domestication occurred during the last century (10, 11). Kiwifruit production and trade in plant material for commercial and breeding purposes has recently increased in Asia, Europe, New Zealand and Chile (12-16), preceding the emergence of disease in some cases by less than a decade.

The first reports of a destructive bacterial canker disease in green-fleshed kiwifruit (*A. chinensis var. deliciosa*) came from Shizuoka, Japan (17, 18). The causal agent was described as *Pseudomonas syringae* pv. *actinidiae* (*Psa*) (18). An outbreak of disease with symptoms similar to those produced by *Psa* was reported to have occurred in 1983-1984 in Hunan, China, though no positive identification was made or isolates stored at that time (17). *Psa* was also isolated from infected green kiwifruit in Korea shortly thereafter (19). The cultivation of more recently developed gold-fruiting cultivars derived from *A. chinensis var. chinensis* (e.g. ‘Hort16A’) began only in the 2000s and an outbreak of global proportions soon followed. The first published notices of the latest outbreak on gold kiwifruit issued from Italy in 2008, with reports from neighbouring European countries, New Zealand, Asia and Chile occurring soon after (1-6, 20). Whole genome sequencing showed the most recent global outbreak of disease was caused by a new lineage of *Psa* (previously referred to as *Psa*-V and now referred to as *Psa*-3), while earlier disease incidents in Japan and Korea were caused by strains forming separate clades referred to *Psa*-1 (previously *Psa*-J) and *Psa*-2 (previously *Psa*-K), respectively (21-24). These clades are marked by substantial variation in their complement of type III secreted effectors, which are required for virulence in *P. syringae*. Despite the surprising level of within-pathovar differences in virulence gene repertoires occurring subsequent to the divergence of these three clades, strains from each clade are capable of infecting and growing to high levels in both *A. chinensis* var. *deliciosa* and *A. chinensis* var. *chinensis* (23).

The severity of the latest global outbreak is largely predicated on the expansion in cultivation of clonally propagated highly susceptible *A. chinensis var. chinensis* cultivars, with trade in plant material and pollen likely providing opportunities for transmission between distant geographic regions. Identifying the source from which *Psa* emerged to cause separate outbreaks remains an important question. Intriguingly, despite the divergence in both the core and flexible genome, these distinct clades nevertheless exhibit evidence of recombination with each other and unknown donors (23). This suggests each lineage of *Psa* emerged from a recombining source population. Definitive evidence for the location, extent of diversity and evolutionary processes operating within this population remain elusive. Early reports suggested China may be the source of the latest global outbreak (22, 23). Although the strains of *Psa* available at that time did not provide unambiguous and well-supported evidence of a Chinese origin, this speculation was based on the fact that kiwifruit are native to China; it is the provenance of the plant material selected for commercial and breeding purposes in China, New Zealand, Italy and other kiwifruit growing regions; there is extensive trade in plant material between all of these regions; and one Chinese isolate was found to carry an integrative and conjugative element (ICE) that was also found in New Zealand *Psa*-3 isolates (22).

In order to strengthen understanding of the population structure, phylogeography and evolutionary dynamics of *Psa*, we isolated *Psa* from cultivated kiwifruit across six provinces in China and obtained additional isolates from South Korea and Japan. Genome sequencing of fifty isolates and the inclusion of an additional thirty previously sequenced isolates show that while China is the origin of the pandemic lineage of *Psa*, only a single clade is currently present in China, while strains from multiple clades are present in both Korea and Japan. Strains from the pandemic lineage are closely related and display reduced pairwise nucleotide diversity relative to other lineages, indicating a more recent origin. Distinct transmission events were responsible for the introduction of the pandemic lineage of *Psa* into New Zealand, Chile and Europe. Two independent transmission events occurred between China and Korea, and two Japanese isolates from 2014 cluster with New Zealand *Psa*. Despite high similarity at the level of the core genome and negligible impact of within-lineage recombination, there has been substantial gene gain and loss even within the single clade from which the global pandemic arose.

## RESULTS

### The phylogeography of *Psa*

The genomes of 50 *P. syringae* pv. *actinidiae* (*Psa*) isolated from symptomatic kiwifruit in China, Korea and New Zealand between 2010 and 2015 were sequenced (Table S1). Combined with 30 *Psa* genomes from earlier outbreaks and different geographic regions (e.g. Italy and Chile), our samples represent the main *Psa* genotypes from the countries producing 90% of kiwifruit production worldwide. The completed reference genome of *Psa* NZ13 (ICMP 18884) comprises a 6,580,291bp chromosome and 74,423bp plasmid (23, 25). Read mapping and variant calling with reference to *Psa* NZ13 chromosome produced a 1,059,722bp nonrecombinant core genome for all 80 genomes, including 2,953 nonrecombinant SNPs. A maximum likelihood phylogenetic analysis showed the four clades of *Psa* known to cause bleeding canker disease in kiwifruit were represented among the 80 strains (Figure 1). The first clade (*Psa*-1) includes the pathotype strain of *Psa* isolated and described during the first recorded epidemic of bleeding canker disease in Japan (1984-1988). The second clade (*Psa*-2) includes isolates from an epidemic in South Korea (1997-1998), and the third clade (*Psa*-3) includes isolates that define the global pandemic lineage (2008-present). A fourth clade (*Psa*-5) is represented by a single strain, as no additional sequences or isolates were available (26). The average between and within-clade pairwise identity is 98.93% and 99.73%, respectively (Table S2). All *Psa* isolated from kiwifruit across six different provinces in China group are members of the same clade: *Psa-3*. A subset of Chinese strains group with the *Psa* isolated during the global outbreak in Italy, Portugal, New Zealand, and Chile. This subset is referred to as the pandemic lineage of *Psa*-3.

**Figure 1.**
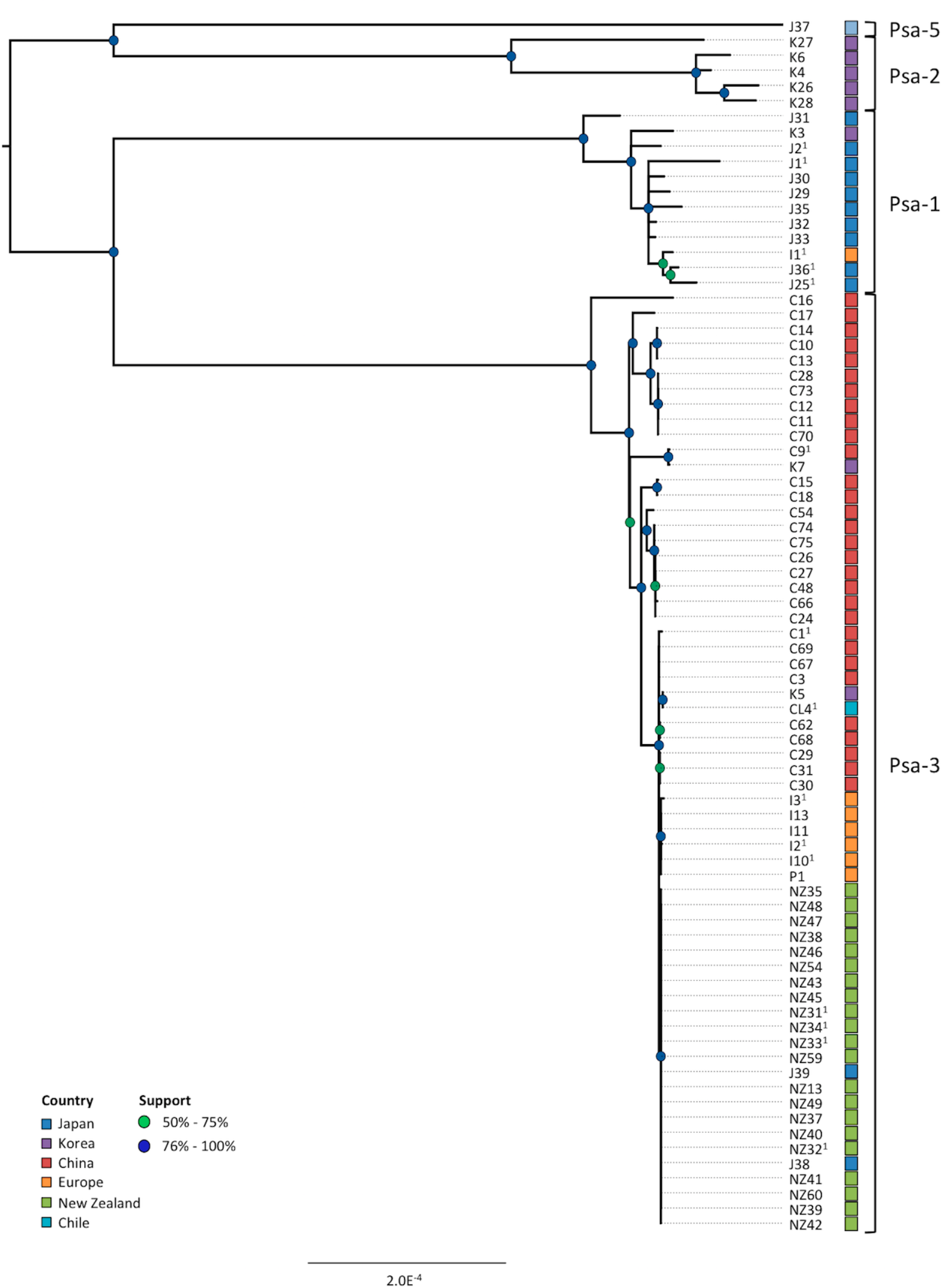
Phylogeny of *Psa*. RaxML Maximum likelihood tree based on 1,059,722bp non-recombinant core genome alignment including 2,953 variant sites. All nodes displayed have bootstrap support values above 50% (50-75% in green, 76-100% in blue). Province of isolation is displayed for Chinese isolates.

In order to obtain greater resolution of the relationships between the new Chinese and pandemic isolates, we identified the 4,853,413bp core genome of all 62 strains in *Psa*-3. The core genome includes both variant and invariant sites and excludes regions either unique to or deleted from one or more strains. To minimize the possibility of recombination affecting the reconstruction of evolutionary relationships and genetic distance within *Psa*-3, ClonalFrameML was employed to identify and remove 258 SNPs with a high probability of being introduced by recombination rather than mutation, retaining 1,948 nonrecombinant SNPs. The within-lineage ratio of recombination to mutation (R/theta) is reduced in *Psa*-3 (6.75 x 10^−2^ ± 3.24 x 10^−5^) relative to between lineage rates (1.27 ± 5.16 x 10^−4^), and the mean divergence of imported DNA within *Psa*-3 is 8.54 x 10^−3^ ± 5.18 x 10^−7^ compared to 5.68 x 10^−3^ ± 1.04 x 10^−8^ between lineages. Although recombination has occurred within *Psa*-3, it is less frequent and has introduced fewer polymorphisms relative to mutation: when accounting for polymorphisms present in recombinant regions identified by ClonalFrameML and/or present on transposons, plasmids, and other mobile elements, more than seven-fold more polymorphisms were introduced by mutation relative to recombination (Table 1). Recombination has a more pronounced impact between clades, where substitutions are slightly more likely to have been introduced by recombination than by mutation (Table 1).

**Table 1.** Origin of SNPs in core genomes

|  |  | **Intergenic** | **Synon** | **Nonsyn** | **Extension** | **Trunc** | **Total** |  |
| --- | --- | --- | --- | --- | --- | --- | --- | --- |
| *Psa-3* | Mutation | 412 | 583 | 927 | 2 | 24 | 1,948 |  |
|  | Recombination | 35 | 136 | 85 | 1 | 1 | 258 | 7.55 |
| **All *Psa*** | Mutation | 457 | 1,493 | 982 | 3 | 18 | 2,953 |  |
|  | Recombination | 355 | 2,212 | 558 | 2 | 5 | 3,132<sup>1</sup> | 0.94 |

<sup>1</sup>For ten positions in the alignment, two recombination events were predicted to occur

### The source of pandemic *Psa*

Data show that there is greater diversity among the Chinese *Psa*-3 population than had been previously identified (Figure 2). Interestingly, clades defining *Psa*-1 and *Psa*-3 exhibit similar levels of diversity (Table S2). These clades share a common ancestor: assuming they are evolving at a similar rate, they may have been present in Japan and China for a similar duration. The strains isolated during the latest pandemic in Italy (I2, I3, I10, I11, I13), Portugal (P1), New Zealand (NZ13, NZ31-35, NZ37-43, NZ45-49, NZ54), Chile (CL4), Japan (J38, J39) and Korea (K5) during the latest kiwifruit canker pandemic cluster with nine Chinese isolates (C1, C3, C29-31, C62, C67-69) (Figure 2). This pandemic lineage exhibits little diversity at the level of the core genome, having undergone clonal expansion only very recently. The NZ isolates form a monophyletic group and share a common ancestor, indicating there was a single transmission event of *Psa* into NZ. Two recently isolated Japanese pandemic *Psa*-3 isolated in 2014 group within the New Zealand isolates, suggesting the pandemic lineage may have been introduced into Japan via New Zealand (Figure 2). Italian and Portuguese pandemic strains also form a separate group, indicative of a single transmission event from China to Italy. China is undoubtedly the source of the strains responsible for the pandemic of kiwifruit canker disease, yet the precise origins of the pandemic subclade remain unclear. Isolates from four different provinces in Western China (Guizhou, Shaanxi, Sichuan and Chongqing) are represented among the pandemic lineage, indicating extensive regional transmission within China after emergence of the pandemic. Yet each province harbouring pandemic isolates also harbors basally diverging *Psa*-3 isolates (Figure 3). With the exception of a group of isolates from Sichuan, there is no phylogeographic signal among the more divergent Chinese strains. This suggests there was extensive regional transmission of *Psa* both prior and subsequent to the emergence of the pandemic subclade in China. Korea harbors both divergent and pandemic subclade *Psa*-3 strains. K5 groups with the Chilean *Psa*-3 strain in the pandemic subclade, while K7 groups with the more divergent Chinese isolates indicating that a transmission event from strains outside the pandemic subclade may have occurred. This pool of diversity therefore represents a reservoir from which novel strains are likely to emerge in the future.

**Figure 2.**
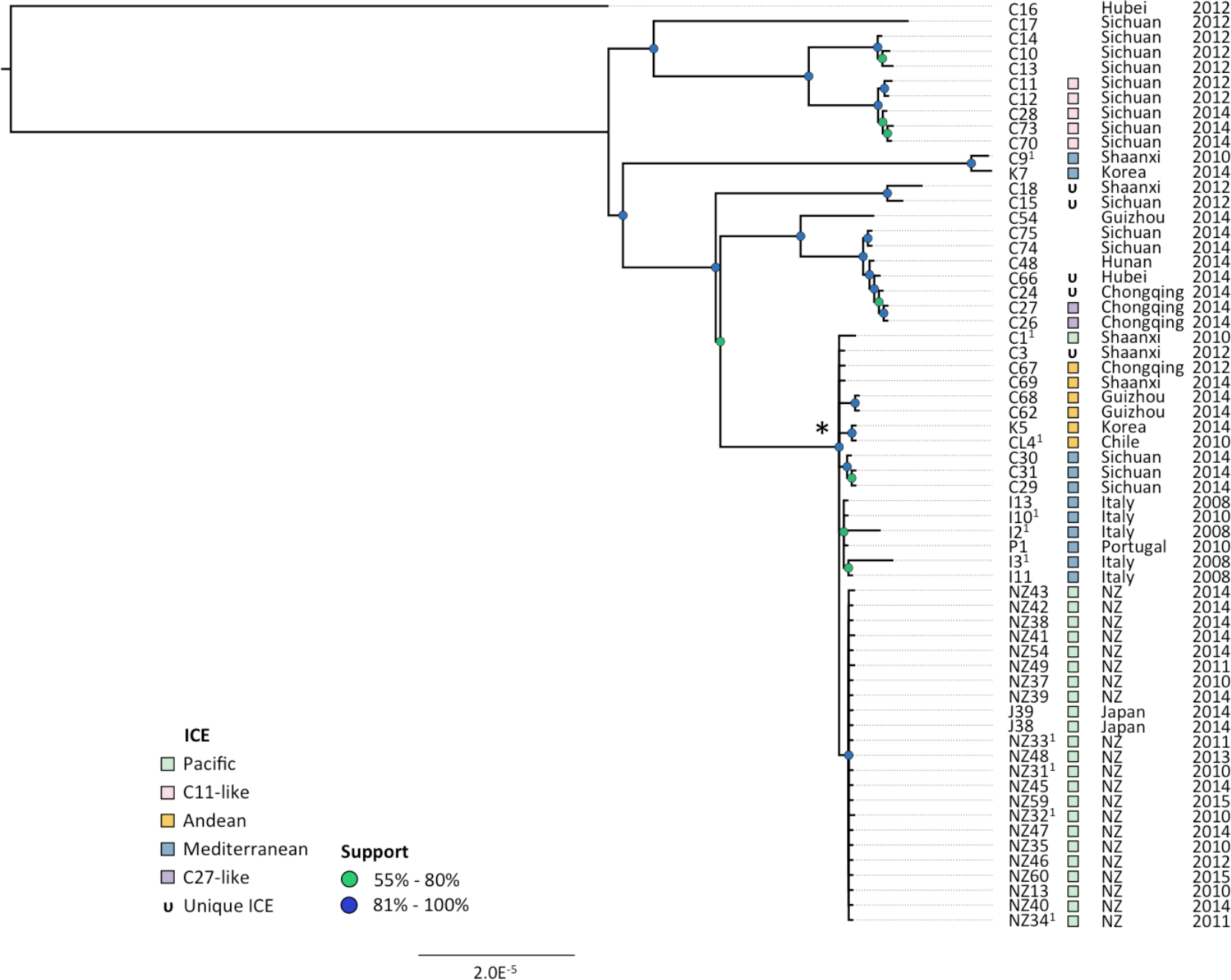
Phylogeny of *Psa*-3. Maximum likelihood tree based on 4,853,155bp non-recombinant core genome alignment including 1,948 variant sites. All nodes displayed have bootstrap support values above 55% (55-80% in green, 81-100% in blue). Year and province (China) or country of isolation is displayed. Integrative and conjugative elements (ICEs) present in each host genome are indicated.

**Figure 3.**
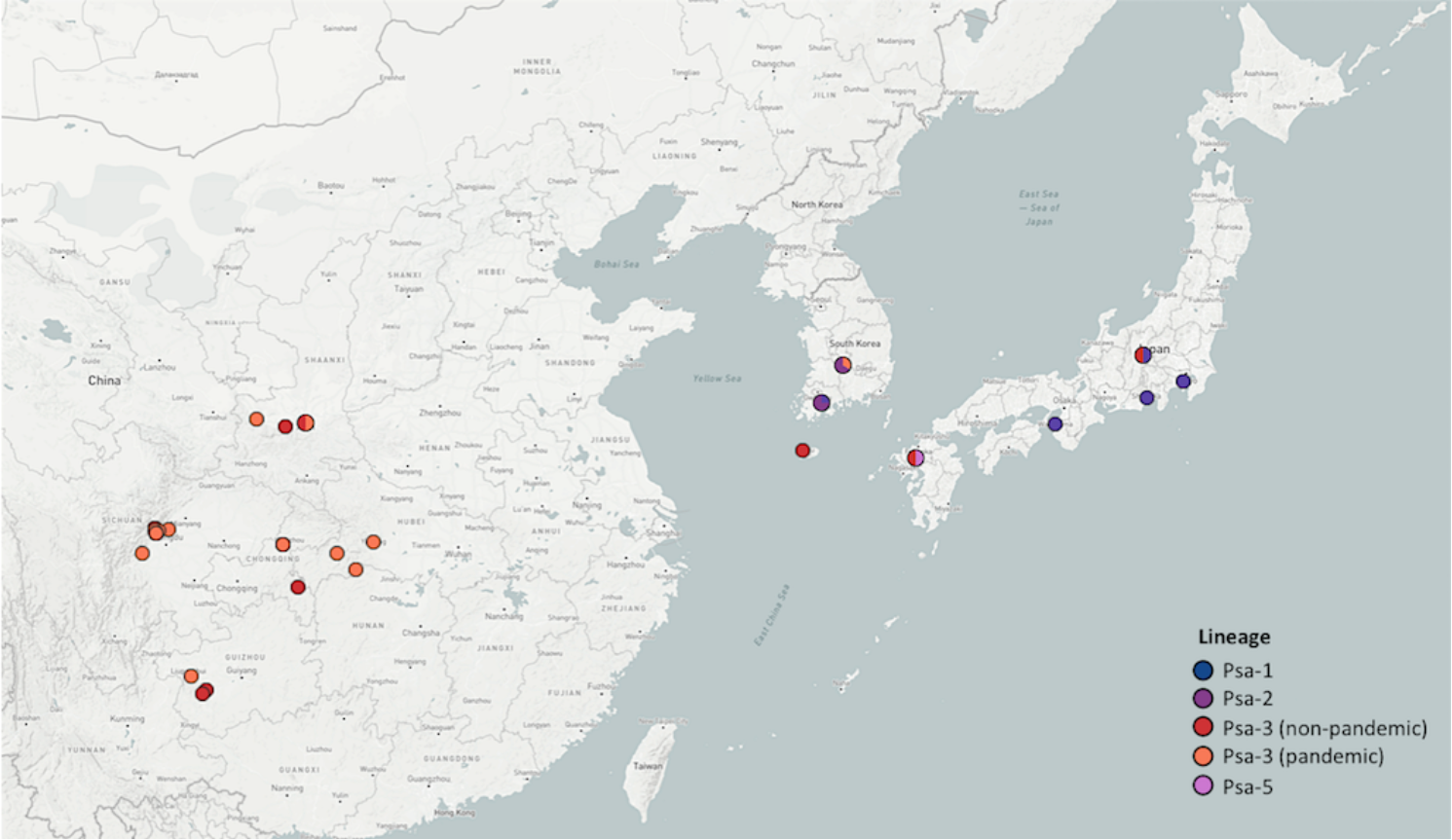
Psa isolation locations in East Asia. Filled circles’ positions correspond to location of isolation (select Japanese and Korean isolates do not have reliable isolation location information). Colour corresponds to the phylogenetic position of the isolates as shown in Figure S1. Map generated in Microreact.

The reduced level of diversity within the core genome of pandemic *Psa*-3 demonstrates these strains have been circulating for a shorter period of time relative to those responsible for earlier outbreaks in both Japan and Korea. In order to estimate the divergence time of the pandemic lineages as well as the age of the most recent common ancestor of all *Psa* clades displaying vascular pathogenicity on kiwifruit, we performed linear regression of root-to-tip distances against sampling dates using the RAxML phylogenies determined from the non-recombinant core genome of all clades and of *Psa*-3 alone. No temporal signal was identified in the data. There were poor correlations between substitution accumulation and sampling dates, indicating the sampling period may have been too short for sufficient substitutions to occur. There may also be variation in the substitution rate within even a single lineage. Forty-four unique non-recombinant SNPs were identified among the 21 pandemic *Psa*-3 genomes sampled over five years in New Zealand (an average of 2.10 per genome) over five years) producing an estimated rate of 8.7 x 10^−8^ substitutions per site per year. The relatively slow substitution rate and the strong bottleneck effect experienced during infections hinders efforts to reconstruct patterns of transmission, as the global dissemination of a pandemic strain may occur extremely rapidly (27, 28). The estimated divergence time of *Psa* broadly considered is likely older than the pandemic and epidemic events with which they are associated: the earliest report of disease cause by lineage 1 occurred in 1984 and the first report of infection from the latest pandemic was issued in 2008.

### Diversification and parallelism among *Psa*-3 isolates

2,206 SNPs mapping to the core genome of *Psa*-3 were identified; 258 of these mapped to recombinant regions identified by ClonalFrameML and/or plasmid, prophage, integrative and conjugative elements, transposons and other mobile genetic elements (Table 1, Figure 4). The highest density of polymorphism occurs in and around the integrative and conjugative element (ICE) in *Psa* NZ13 (Figure 4). Of the 1,948 SNPs mapping to the non-recombinant non-mobile core genome, 58.1% (1,132) are strain specific. Most strain-specific SNPs are found in the two most divergent members of the lineage: *Psa* C16 and C17, with 736 and 157 strain-specific SNPs, respectively. The remaining isolates have an average of 4.0 strain-specific SNPs, ranging from 0 to 44 SNPs per strains. There are 816 SNPs shared between two or more *Psa*-3 strains. The pandemic clade differs from the more divergent Chinese strains by 72 shared SNPs. Within the pandemic lineage there are 125 strain-specific SNPs, an average of 3.1 unique SNPs per strain (ranging from 0-27 SNPs) and an additional 29 SNPs shared among pandemic strains. Protein-coding sequence accounts for 88.4% of the non-recombinant, gap-free core genome of this clade. We observed that 78.9% (1,536/1,948) of mutations occurred in protein coding sequence, significantly different from the expectation (1,722/1,948) in the absence of selection (Pearson’s χ^2^ test: *P* < 0.0001, χ^2^ = 173.46). This suggests there is selection against mutations occurring in protein coding sequences. Of the 953 substitutions introduced by mutation in *Psa*-3, 927 resulted in amino acid substitutions, two resulted in extensions and 24 resulted in premature truncations.

**Figure 4.**
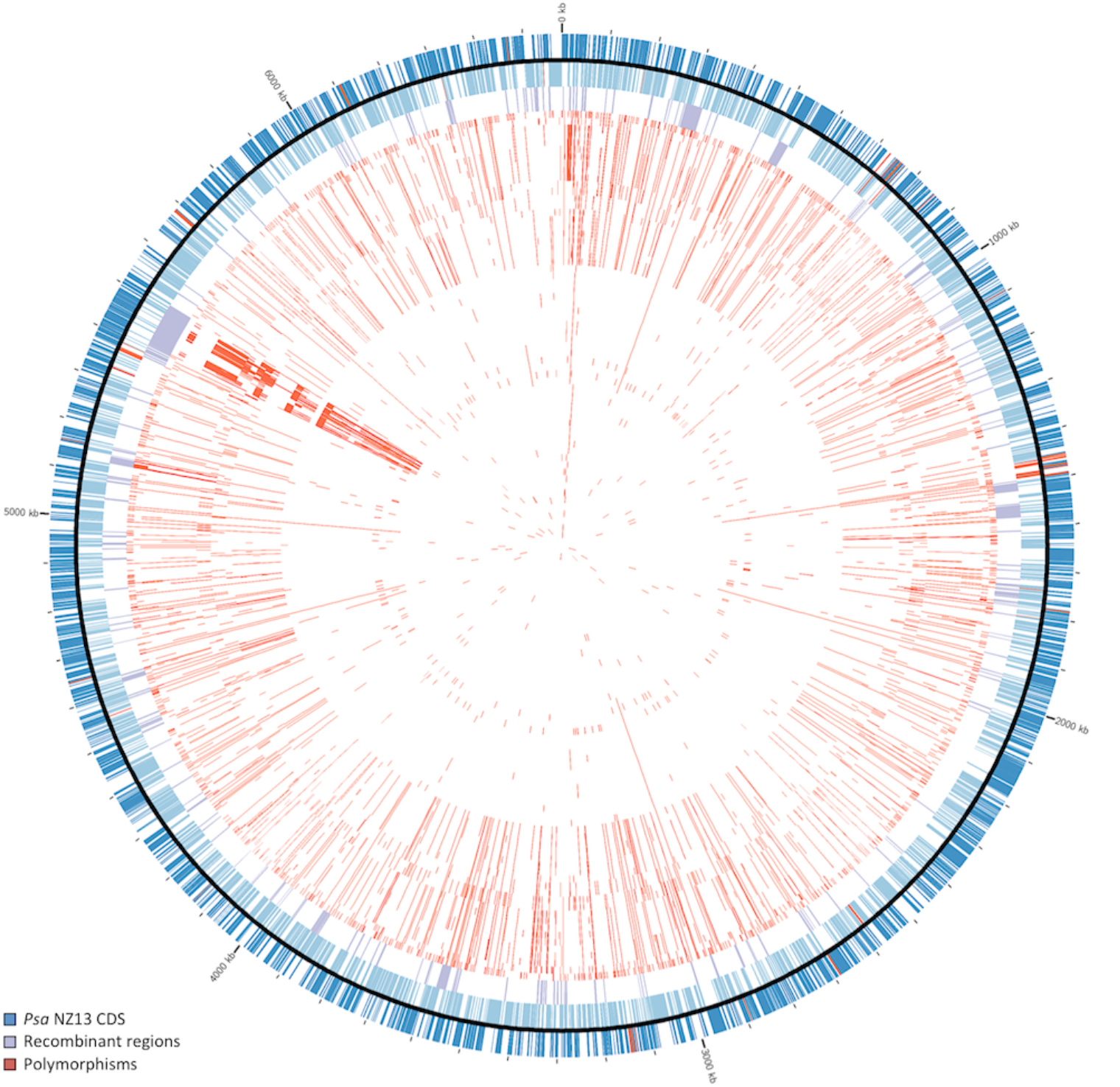
Genomic context of polymorphisms in *Psa*-3. Polymorphisms and recombinant regions mapped onto *Psa* NZ13 reference genome using CIRCOS (64). *Psa* NZ13 CDS are displayed in the first and second ring (blue), with annotated Type 3 secretion system and effectors highlighted (red). Inner rings display polymorphisms in *Psa*-3 genomes ordered from most to least divergent relative to *Psa* NZ13 (see Figure 2). The most polymorphic region corresponds to the location of the integrative and conjugative element (ICE) in *Psa* NZ13.

Multiple synonymous and non-synonymous mutations were identified in 269 genes. The accumulation of multiple independent mutations in the same gene may be a function of gene length, mutational hotspots or directional selection-A range of hypothetical proteins, membrane proteins, transporters, porins and type III and IV secretion system proteins acquired between two and seven mutations. The fitness impact of these mutations - and the 38 amino-acid changing mutations in the ancestor of the pandemic subclade – is unknown, yet it is possible these patterns are the outcome of selective pressures imposed during bacterial residence within a similar host niche.

Two substitutions are shared exclusively by the European pandemic strains (AKT28710.1 G1150A and AKT33438.1 T651C) and one silent substitution in a gene encoding an acyltransferase superfamily protein (AKT31915.1 C273T) is shared among the European pandemic and six of nine Chinese pandemic strains (C3, C29-31, C67, C69). As these six Chinese pandemic strains were isolated from Shaanxi, Sichuan and Chongqing, they do not provide any insight into the precise geographic origins of the European pandemic *Psa*-3, though transmission from China to Italy is likely concomitant with dissemination of the pandemic lineage across China. Six conserved and diagnostic polymorphisms are present in the pandemic New Zealand and Japanese isolates (Table S3). One of these is a silent substitution in an ion channel protein (AKT31947.1 A213G), another is an intergenic (T->G) mutation at position 362,522 of the reference *Psa* NZ13 chromosome and the remaining four are nonsynonymous substitutions in an adenylyltransferase (AKT32845.1, W977R); chromosome segregation protein (AKT30494.1, H694Q); cytidylate kinase (AKT29651.1, V173L) and peptidase protein (AKT32264.1, M418K).

The type III secretion system is known to be required for virulence in *P. syringae*. A 44,620bp deletion event in *Psa* C17 resulted in the loss of 42 genes encoding the structural apparatus and conserved type III secreted effectors in *Psa* C17. This strain is highly compromised in its ability to grow in *A. chinensis* var. *deliciosa* ‘Hayward’, attaining 1.2 x 10^7^ cfu/g three days post inoculation (dpi) and declining to 8.8 x 10^4^ cfu/g at fourteen dpi (Figure S1). This is a marked reduction compared to *Psa* NZ13, which attains 3.0 x 10^9^ and 4.2 x 10^7^ cfu/g three and fourteen dpi, respectively. *Psa* C17 nevertheless multiplies between day 0 and day 3, indicating that even in the absence of type III-mediated host defense disruption, *Psa* may still proliferate in host tissues. The loss of the TTSS does not inhibit the growth of *Psa* C17 as strongly in the more susceptible *A. chinensis* var. *chinensis* ‘Hort16A’ cultivar.

Two potentially significant deletion events occurred in the ancestor of the pandemic subclade: a frameshift caused by a mutation and single base pair deletion in a glucan succinyltransferase (*opgC*) and a 6,456bp deletion in the *wss* operon (Figure S2). Osmoregulated periplasmic glucans (OPGs, in particular *opgG* and *opgH*) are required for motility, biofilm formation and virulence in various plant pathogenic bacteria and fungi (29-31). Homologs of *opgGH* remain intact in the pandemic subcclade, yet the premature stop mutation in *opgC* likely results in the loss of glucan succinylation. The soft-rot pathogen *Dickeya dadantii* expresses OpgC in high osmolarity conditions, resulting in the substitution of OPGs by O-succinyl residues (32). *D. dadantii opgC* deletion mutants did not display any reduction in virulence (32). *Psa* is likely to encounter high osmolarity during growth and transport in xylem conductive tissues, yet the impact of the loss of *opgC* on *Psa* fitness has yet to be determined. The most striking difference between the pandemic subclade and more divergent Chinese *Psa*-3 strains is the deletion of multiple genes involved in cellulose production and acetylation of the polymer (Figure S2) (33). The loss of cellulose production and biofilm production is not associated with a reduction in growth or symptom development of *P. syringae* pv. *tomato* DC3000 on tomato, but may enhance bacterial spread through xylem tissues during vascular infections (34). In *P. fluorescens* SBW25 deletion of the Wss operon significantly compromises ability to colonise plant surfaces and in particular the phyllosphere of sugar beet (*Beta vulgaris*) seedlings (35). It is possible that loss of this locus aids movement through the vascular system and / or dissemination between plants, by limiting capacity for surface colonization and biofilm formation.

### Dynamic genome evolution of *Psa*-3

Despite the high similarity within the core genome, extensive variation is evident in the pangenome of *Psa*-3 (Figure 5). The core genome (4,339 genes in 99-100% of strains, and 674 genes in 95%-99% of strains, or 58-62 genomes) comprises 50.5% of the total pangenome (9,931 genes). 968 genes are present in 15-95% of strains (9-57 genomes), the so-called ‘shell genes’ (Figure 5). The flexible genome is comprised of the ‘shell’ and ‘cloud’ genes; the latter describes genes present in 0-15% of strains (one to six genomes in this case). Cloud genes contribute most to the flexible genome: 3,950 genes are present in one to six strains. This is a striking amount of variation in a pathogen described as clonal and monomorphic. It should be noted that sequencing and assembly quality will impact annotation and pangenome estimates: omitting the low quality J39 assembly results in a core and soft-core genome differing by 18 genes and a reduction of the cloud by 275. Despite a relatively slow rate of mutation and limited within-clade homologous recombination, the amount of heterologous recombination demonstrates that the genomes of these pathogens are highly labile. Mobile genetic elements like bacteriophage, transposons and integrases make a dramatic contribution to the flexible genome. Integrative and conjugative elements (ICEs) are highly mobile elements and have recently been demonstrated to be involved in the transfer of copper resistance in *Psa* (36). Prodigious capacity for lateral gene transfer creates extreme discordance between ICE type, host phylogeny and host geography making these regions unsuitable markers of host evolution and origin.

**Figure 5.**
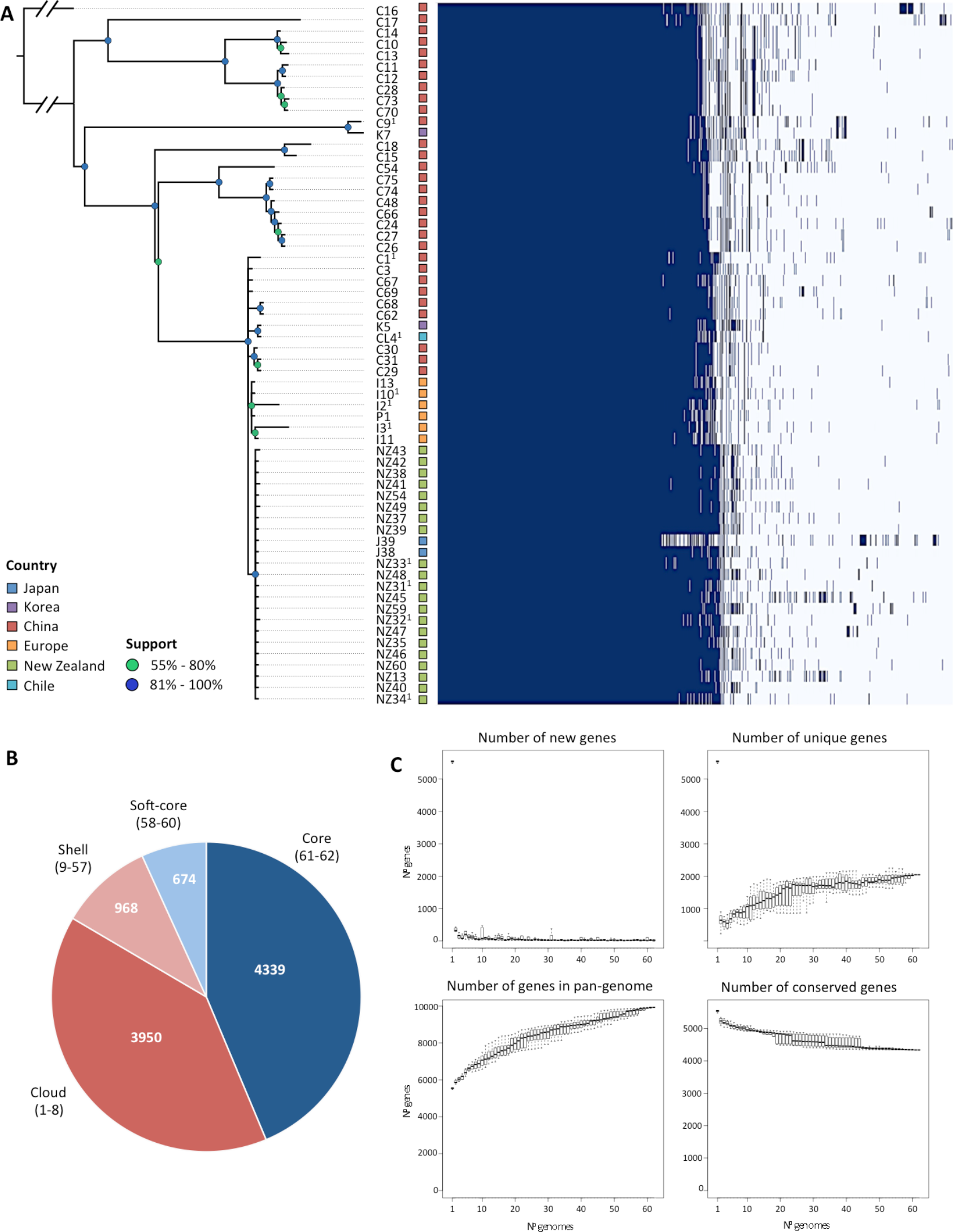
Pangenome of *Psa*-3. A. Presence/absence matrix of all core and accessory genes in *Psa*-3, ordered according to strains’ phylogenetic relationships. Country of isolation is indicated at left. B. The core and flexible genome of *Psa*-3. The core, soft-core, shell and cloud genomes are defined according to the numbers in parentheses. C. Number of new, unique and conserved genes with addition of each genome.

Three divergent ICEs have been previously described from the global pandemic lineage (23). Within *Psa*-3 ICEs were found in 53 of 62 isolates (nine of the divergent Chinese isolates were devoid of any such element) (Figure 2). No phylogeographic signal is evident. For example, strains from Sichuan, Shaanxi, Korea, Italy and Portugal share an identical ICE. Even within a single Chinese province, mulitple ICEs exist (Shaanxi and Sichuan isolates harbour four and three different ICEs, respectively). Moreover, ICE host range is not limited to *Psa* alone: the ICE found in every NZ isolate (and also recorded in Chinese isolate C1) exists in essentially identical form in a strain of *P. syringae* pv. *avellenae* CRAPAV013 isolated from hazelnut in 1991 in Latina, Italy (it exhibits 98% pairwise identity, differing from the New Zealand ICE by a transposon, 66bp deletion and a mere 6 SNPs).

## DISCUSSION

We have described an endemic population of *Psa* infecting cultivated kiwifruit in China. All *Psa* isolated within China are members of the same lineage as that responsible for the latest pandemic. The pandemic strains isolated in Italy, Portugal, Chile and New Zealand form a subclade within this lineage along with a subset of Chinese isolates, indicating that the pandemic ultimately emerged from the Chinese population of lineage 3 strains. Italian pandemic strains share a SNP with six of nine Chinese pandemic *Psa* strains, indicating there was likely a direct transmission event from China to Italy prior to 2008. The New Zealand isolates share six clade-defining mutations, indicating that a separate and single transmission event was responsible for the outbreak of disease there. Identification of the transmission pathway introducing *Psa* into New Zealand is dependent on obtaining a sample of *Psa* sharing some or all of the mutations characteristic of NZ *Psa* from either the overseas source population or from infected plant material arriving into New Zealand from an overseas location. The relatively low mutation rate in the core genome of *Psa* places a lower boundary on the ability of genomic epidemiology to resolve transmission events occurring either rapidly (as a consequence of human-mediated long-distance dissemination) or at a local scale. The Japanese pandemic strains cluster with the NZ strains, and share all six clade-defining mutations. This suggests that pandemic *Psa*-3 was either introduced into Japan via New Zealand, or from the same as-yet unknown region in China from which transmission to New Zealand occurred. *Psa*-3 was first identified as causing disease in four prefectures across Japan in April 2014 (5). Japan imported pollen and plant material from both China and New Zealand prior and subsequent to *Psa*-3 detection in both those countries, though the amount of pollen imported from New Zealand in 2012 (349 kg) and 2013 (190 kg) far outweighed the amount imported from China (1 kg in both 2012 and 2013) (37).

Our phylogeographic study of a single lineage giving rise to a pandemic in *P. syringae* has revealed far greater diversity than was previously appreciated. Extensive diversity between *Psa* isolates collected from *Actinidia* spp. was observed in the same province. The amount of diversity present within lineage 3 indicates this population was present and circulating in China before the pandemic began. The emergence of the pandemic subclade moreover has not resulted in the displacement of more ancestral strains: both pandemic and divergent lineage 3 *Psa* were isolated from four out of six provinces.

Strains from three different clades have been isolated in both Korea and Japan, while China harbours strains from only a single clade. The most basal clades of canker-causing *Psa* are comprised of Korean strains isolated between 1997 and 2014 (*Psa*-2) and a member of the recently identified lineage *Psa*-5. One early isolate (*Psa* K3, 1997) groups with the Japanese isolates in lineage 1, and a more recent Korean isolate *Psa* K7 (2014) groups with the more diverse Chinese isolates in *Psa*-3. Korea therefore harbours a more diverse population of *Psa* than China, with strains from three distinct lineages of *Psa* (1, 2 and non-pandemic subclade 3). The novel group of *Psa* recently identified in Japan (*Psa*-5) appears to share an ancestor with the Korean *Psa*-2 strains. With the recent dissemination of pandemic *Psa*-3 and the historical presence of *Psa*-1 in Japan, the Japanese population of *Psa* is comprised of three distinct clades of *Psa* (1, 5 and pandemic *Psa*-3). Though no strains from *Psa*-1 have been isolated in either Japan or Korea since 1997, at least two lineages currently coexist in both Japan and Korea. This strongly suggests that the source population of all *Psa* is not China, but likely resides in either Korea or Japan. The potential transmission of a non-pandemic *Psa*-3 strain from China to Korea and the identification of a new clade in Japan supports our earlier assertion that variants will continue to emerge to cause local epidemics and global pandemics in the future (23).

Considering that the divergence time of this monophyletic pathovar predates the commercialisation of kiwifruit by hundreds if not thousands of years, *Psa* is likely associated with a non-domesticated host(s) in the wild. Both *A. chinensis var. deliciosa* or *A. chinensis var. chinensis* are found in natural ecosystems and have overlapping habitat ranges with cultivated kiwifruit in many areas. However, despite isolating 746 *Pseudomonas* strains from both wild and cultivated kiwifruit during this sampling program, we did not identify *Psa* among any of the 188 *Pseudomonas* spp. isolated from 98 wild *A. chinensis var. deliciosa* or *A. chinensis var. chinensis* sampled across China (Table S4). Very few *Actinidia* spp. have ranges extending to South Korea and Japan: *A. arguta, A. kolomikta, A. polygama* and *A. rufa. A. arguta* are broadly distributed across both Korea and Japan. Early work by Ushiyama *et al*. (1992) found that *Psa* could be isolated from symptomatic *A. arguta* plants in Japan. The possibility that this wild relative of kiwifruit harbours diverse strains of *Psa* that may emerge to cause future outbreaks is currently under investigation. Alternately, a host shift from another domesticated crop may have occurred after expansion in kiwifruit cultivation.

Numerous epidemiological studies of human pathogens have demonstrated environmental or zoonotic origins, but there are few such studies of plant pathogens (23, 38-48). Where ecological and genetic factors restrict pathogens to a small number of plant hosts some progress has been made, but for facultative pathogens such as *P. syringae* that colonise multiple hosts and are widely distributed among both plant and non-plant habitats, the environmental reservoirs of disease and factors affecting their evolutionary emergence are difficult to unravel (49, 50).

The emergence of *Psa* over the last three decades - concomitant with domestication of kiwifruit-offers a rare opportunity to understand the relationship between wild populations of both plants and microbes and the ecological and evolutionary factors driving the origins of disease, including the role of agriculture. It is now possible to exclude China as the native home to the source population, but the precise location remains unclear. Nonetheless, it is likely, given the extent of diversity among *Psa* isolates and the time-line to domestication, that ancestral populations exist in non-agricultural plant communities. Attention now turns to Korea and Japan and in particular the interplay between genetic and ecological factors that have shaped *Psa* evolution.

## MATERIALS AND METHODS

### Bacterial strains and sequencing

Samples were procured by isolation from symptomatic plant tissue. Bacterial strain isolations were performed from same-day sampled leaf and stem tissue by homogenising leaf or stem tissue in 800uL 10mM MgSO_4_ and plating the homogenate on *Pseudomonas* selective media (King’s B supplemented with cetrimide, fucidin and cephalosporin, Oxoid). Plates were incubated 48 hours between 25 and 30°C. Single colonies were restreaked and tested for oxidase activity, and used to inoculate liquid overnight cultures in KB. Strains were then stored at -80°C in 15% glycerol and the remainder of the liquid culture was reserved for genomic DNA isolation by freezing the pelleted bacterial cells at -20°C. Genomic DNA extractions were performed using Promega Wizard 96-well genomic DNA purification system.

Initial strain identification was performed by sequencing the citrate synthase gene (*cts*, aka *gltA* (51)). Subsequent to strain identification, paired-end sequencing was performed using the Illumina HiSeq 2500 platform (Novogene, Guangzhou, China). Additional paired-end sequencing was performed at New Zealand Genomics Limited (Auckland, New Zealand) using the MiSeq platform, and raw sequence reads from some previously published isolates were shared by Mazzaglia *et al*. (24).

### Variant Calling and Recombination Analyses

The completely sequenced genome of *Pseudomonas syringae* pv. *actinidiae* NZ13 was used as a reference for variant calling. A near complete version of this genome was used as a reference in our previous publication and subsequently finished by Templeton *et al*. (2015), where it is referred to as ICMP18884 (23, 25). Variant calling was performed on all *P. syringae* pv. *actinidiae* isolates for which read data was available.

Read data was corrected using the SPADEs correction module and Illumina adapter sequences were removed with Trimmomatic allowing 2 seed mismatches, with a palindrome and simple clip threshold of 30 and 10, respectively (52, 53). Quality-based trimming was also performed using a sliding window approach to clip the first 10 bases of each read as well as leading and trailing bases with quality scores under 20, filtering out all reads with a length under 50 (53). PhiX and other common sequence contaminants were filtered out using the Univec Database and duplicate reads were removed (54).

Reads were mapped to the complete reference genome *Psa* NZ13 with Bowtie2 and duplicates removed with SAM Tools (55, 56). Freebayes was used to call variants with a minimum base quality 20 and minimum mapping quality 30 (57). Variants were retained if they had a minimum alternate allele count of 10 reads and fraction of 95% of reads supporting the alternate call. The average coverage was calculated with SAM Tools and used as a guide to exclude overrepresented SNPs (defined here as threefold higher coverage than the average) which may be caused by mapping to repetitive regions. BCFtools filtering and masking was used to generate final reference alignments including SNPs falling within the quality and coverage thresholds described above and excluding SNPs within 3bp of an insertion or deletion (indel) event or indels separated by 2 or fewer base pairs. Invariant sites with a minimum coverage of 10 reads were also retained in the alignment, areas of low (less than 10 reads) or no coverage are represented as gaps relative to the reference.

Freebayes variant calling includes indels and multiple nucleotide insertions as well as single nucleotide insertions, however only SNPs were retained for downstream phylogenetic analyses. An implementation of ClonalFrame suitable for use with whole genomes was employed to identify recombinant regions using a maximum likelihood starting tree generated by RaxML (58, 59). All substitutions occurring within regions identified as being introduced due to recombination by ClonalFrameML were removed from the alignments. The reference alignments were manually curated to exclude substitutions in positions mapping to mobile elements such as plasmids, integrative and conjugative elements and transposons.

### Phylogenetic Analysis

The maximum likelihood phylogenetic tree of 80 *Psa* strains comprising new Chinese isolates and strains reflecting the diversity of all known lineages was built with RAxML (version 7.2.8) using a 1,062,844bp core genome alignment excluding all positions for which one or more genomes lacked coverage of 10 reads or higher (59). Removal of 3,122 recombinant positions produced a 1,059,722bp core genome alignment including 2,953 variant sites. Membership within each phylogenetic clade corresponds to a minimum average nucleotide identity of 99.70%. The average nucleotide identity was determined using a BLAST-based approach in JspeciesWS (ANIb), using a subset of 32 *Psa* genome assemblies spanning all clades (60). In order to fully resolve the relationships between more closely related recent outbreak strains, a phylogeny was constructed using only the 62 *Psa*-3 strains. This was determined using a 4,853,155bp core genome alignment (excluding 258 recombinant SNPs), comprising invariant sites and 1,948 non-recombinant SNPs and invariant sites. Trees were built with the generalized time-reversible model and gamma distribution of site-specific rate variation (GTR+Γ) and 100 bootstrap replicates. *Psa* C16 was used to root the tree as this was shown to be the most divergent member of the phylogeny when including strains from multiple lineages. Nodes shown have minimum bootstrap support values of 50.

### Identification of the core and mobile genome

Genomes were assembled with SPAdes using the filtered, trimmed and corrected reads (52). Assembly quality was improved with Pilon and annotated with Prokka (61, 62). The pangenome of *Psa*-3 was calculated using the ROARY pipeline (63). Orthologs present in 61 (out of a total of 62) genomes were considered core; presence in 58-60, 9-57 and 1-8 were considered soft-core, shell and cloud genomes, respectively. BLAST-based confirmation was used to confirm the identity predicted virulence or pandemic-clade-restricted genes in genome assemblies.

### Pathogenicity assays

Growth assays were performed using both stab inoculation as in McCann *et al*. (2013) an initial inoculum of 10^8^cfu/mL and four replicate plants at day 0 and six at all subsequent sampling time points. Bacterial density in inoculated tissue was assessed by serial dilution plating of homogenized tissue. Statistical significance between each treatment at each time point was assessed using two-tailed t-tests with uneven variance.

## ACKNOWLEDGEMENTS

We gratefully acknowledge the assistance of the following guides, teachers, and graduate assistants who helped us identify sample locations in China: Junjie Gong, Yancang Wang, Shengju Zhang, Zupeng Wang, Yangtao Guo, Meiyan Chen, Kuntong Li, Moucai Wang, Jiaming He, Yonglin Zhao, Zhongshu Yu, Yan Lv, Mingfei Yao, Shihua Pu, Tingwen Huang, Qiuling Hu, Caizhi He, and Jiaqing Peng. Derk Wachsmuth at Max Planck Institute for computing server support. James Connell for assistance with biosecurity regulations. Members of the Rainey and Huang labs for discussion. Joel Vanneste for contributing strains. This work was funded by grants from the New Zealand Ministry for Business, Innovation and Employment (C11X1205), Canada Natural Sciences and Engineering Research Council (NSERC PDF), Chinese Academy of Sciences President’s International Fellowship Initiative (Grant NO. 2015PB063), China Scholarship Council (Grant NO. 201504910013), National Natural Science Foundation of China (Grant NO. 31572092), Science and Technology Service Network Initiative Foundation of The Chinese Academy of Sciences (Grant NO. KFJ-EW-STS-076), Protection and utilization of Crop Germplasm Resources Foundation of Ministry of Agriculture (Grant NO. 2015NWB027).

---

**Table S1.** Strains

| ID | WGS origin | Host plant | Country | Year | Other collection/alias | Contigs | N50 |
| --- | --- | --- | --- | --- | --- | --- | --- |
| C1<sup>1</sup> | Mazzaglia *et al.* (2012) | *A. chinensis* 'Hongyang' | China, Shaanxi, Wei | 2010 | CH2010-6, M7 | 470 | 24,560 |
| C3 | This paper | *A. deliciosa* 'Hayward' | China, Shaanxi, Xi'an, Zhouzhi | 2012 | ZY2, CC770 | 318 | 46,356 |
| C9<sup>1</sup> | Butler *et al.* (2013) |  | China, Shaanxi, Wei County | 2010 | M228 | 346 | 38,960 |
| C10 | This paper | *A. chinensis* 'Hongyang' | China, Sichuan, Shifang | 2012 | 850.1.1. CC822 | 325 | 45,191 |
| C11 | This paper | *A. chinensis* 'Hongyang' | China, Sichuan, Shifang | 2012 | 850.2.2. CC823 | 335 | 50,545 |
| C12 | This paper | *A. chinensis* 'Hongyang' | China, Sichuan, Shifang | 2012 | 850.4.1. CC826 | 339 | 48,835 |
| C13 | This paper | *A. chinensis* 'Hongyang' | China, Sichuan, Shifang | 2012 | 850.5.1. CC827 | 326 | 44,835 |
| C14 | This paper | *A. chinensis* 'Hongyang' | China, Sichuan, Shifang | 2012 | 850.6.1. CC828 | 326 | 44,683 |
| C15 | This paper | *A. chinensis* 'Hort16A' | China, Sichuan, Pengzhou, Cifeng | 2012 | 913.1.1. CC835 | 327 | 54,039 |
| C16 | This paper | *A. chinensis* 'Hongyang' | China, Hubei, Enshi, Jianshi | 2012 | 913.5.1, CC836 | 386 | 42,362 |
| C17 | This paper | *A. chinensis* 'Hongyang' | China, Sichuan, Qionglai, Huojing | 2012 | 913.10.1, CC837 | 366 | 43,680 |
| C18 | This paper | *Actinidia* sp. | China, Shaanxi, Baoji, Meixian | 2012 | 913.15.1, CC838 | 325 | 50,834 |
| C24 | This paper | *A. chinensis* 'Hongyang' | China, Chongqing, Wanzhou, Houshan | 2014 | 120L3 | 338 | 48,742 |
| C26 | This paper | *A. chinensis* 'Hongyang' | China, Chongqing, Wanzhou, Houshan | 2014 | 124L1 | 343 | 47,302 |
| C27 | This paper | *A. chinensis* 'Hongyang' | China, Chongqing, Wanzhou, Houshan | 2014 | 124L7 | 364 | 48,757 |
| C28 | This paper | *A. chinensis* 'Hongyang' | China, Sichuan, Dujiangyan, Xujia | 2014 | 139S2 | 343 | 46,114 |
| C29 | This paper | *A. deliciosa* 'Hayward' | China, Sichuan, Dujiangyan, Hongkou | 2014 | 163S1 | 321 | 41,599 |
| C30 | This paper | *A. chinensis* | China, Sichuan, Dujiangyan, Hongkou | 2014 | 165L4 | 334 | 43,004 |
| C31 | This paper | *A. chinensis* | China, Sichuan, Dujiangyan, Hongkou | 2014 | 166L2 | 303 | 43,291 |
| C48 | This paper | *A. chinensis* | China, Hunan, Changde City, Shimen | 2014 | 50L1 | 351 | 47,058 |
| C54 | This paper | *A. deliciosa* | China, Guizhou, Liupanshui, Panxian | 2014 | 77L5 | 360 | 47,421 |
| C62 | This paper | *A. chinensis* 'Jinyan' | China, Guizhou, Liupanshui, Liuzhi | 2014 | GZ410 | 327 | 45,829 |
| C66 | This paper | *A. chinensis* 'Hongyang' | China, Hubei, Yichang, Yiling | 2014 | YC5 | 333 | 48,710 |
| C67 | This paper | *A. chinensis* 'Hongyang' | China, Chongqing, Qianjiang, Jinxi | 2012 | 163W4 | 310 | 46,111 |
| C68 | This paper | *A. chinensis* 'Jinyan' | China, Guizhou, Liupanshui, Liuzhi | 2014 | GZ3-5 | 347 | 45,058 |
| C69 | This paper | *A. chinensis* 'Jinyan' | China, Shaanxi, Xi'an, Zhouzhi | 2014 | SH1-14 | 304 | 47,863 |
| C70 | This paper | *A. chinensis* | China, Sichuan, Dujiangyan, Xujia | 2014 | 141S5 | 327 | 48,715 |
| C73 | This paper | *A. chinensis* 'Hongyang' | China, Sichuan, Dujiangyan, Xujia | 2014 | 139L10 | 365 | 48,937 |
| C74 | This paper | *A. chinensis* 'Hongyang' | China, Sichuan, Dujiangyan, Xiangge | 2014 | 148L1 | 326 | 47,058 |
| C75 | This paper | *A. chinensis* 'Hongyang' | China, Sichuan, Dujiangyan, Xiangge | 2014 | 148L4 | 336 | 47,058 |
| CL4<sup>1</sup> | Butler *et al.* (2013) | *A. deliciosa* | Chile, Maule | 2010 |  | 370 | 37,470 |
| I1<sup>1</sup> | Marceletti *et al.* (2011) | *A. deliciosa* 'Hayward' | Italy, Roma | 1992 | NCPB3871 | 405 | 27,730 |
| I2<sup>1</sup> | Marceletti *et al.* (2011) | *A. chinensis* 'Hort16A' | Italy, Latina | 2008 | CRAFRU8.43 | 523 | 22,372 |
| I3<sup>1</sup> | Mazzaglia *et al.* (2012) | *A. chinensis* 'Hort16A' | Italy, Lazio | 2008 | CFBP 7286 | 329 | 31,420 |
| I10<sup>1</sup> | Butler *et al.* (2013) | *A. deliciosa* | Italy, Roma | 2010 | ICMP18744,<br>CRAFRU11.41 | 358 | 35,904 |
| I11 | Mazzaglia *et al.* (2012) | *A. chinensis* 'Jin Tao' | Italy, Veneto | 2008 | CFBP 7285 | 359 | 33,568 |
| I13 | Mazzaglia *et al.* (2012) | *A. deliciosa* 'Hayward' | Italy, Lazio | 2008 | CFBP 7287 | 357 | 36,668 |
| J1<sup>1</sup> | Baltrus *et al.* (2011) | *A. deliciosa* | Japan, Kanagawa | 1984 | MAFF 302091 | 248 | 65,551 |
| J2<sup>1</sup> | Mazzaglia *et al.* (2012) | *A. chinensis* | Japan | 1988 | PA459 | 634 | 17,643 |
| J25<sup>1</sup> | Mazzaglia *et al.* (2012) | *A. deliciosa* 'Hayward' | Japan, Shizuoka | 1984 | KW41 | 570 | 18,393 |
| J29 | McCann *et al.* (2013) | *A. arguta* | Japan, Kanagawa | 1987 | MAFF302133, JpSar1 | 412 | 30,107 |
| J30 | McCann *et al.* (2013) | *A. arguta* | Japan, Kanagawa | 1987 | MAFF302134, JpSar2 | 401 | 32,586 |
| J31 | McCann *et al.* (2013) | *A. deliciosa* 'Hayward' | Japan, Kanagawa | 1987 | MAFF302143, JpKiw4 | 1723 | 4,505 |
| J32 | McCann *et al.* (2013) | *A. deliciosa* 'Hayward' | Japan, Wakayama | 1988 | MAFF302145, JpWa1 | 465 | 33,908 |
| J33 | McCann *et al.* (2013) | *A. deliciosa* 'Hayward' | Japan, Wakayama | 1988 | MAFF302146, JpWa2 | 410 | 31,305 |
| J35 | McCann *et al.* (2013) | *A. deliciosa* 'Hayward' | Japan, Shizuoka | 1984 | NCPB 3739, Kw11 | 368 | 39,207 |
| J36<sup>1</sup> | Butler *et al.* (2013) | *A. deliciosa* 'Hayward' | Japan, Shizuoka | 1984 | Kw1 | 417 | 27,018 |
| J37<sup>1</sup> | Fujikawa & Sawada (2015) |  | Japan, Saga | 2014 | PRJDB2950 | 291 | 50,639 |
| J38 | This paper | *A. chinensis* 'Hort16A' | Japan | 2014 |  | 313 | 48,009 |
| J39<sup>2</sup> | This paper | *A. chinensis* 'Hort16A' | Japan, Saga | 2014 |  | 1221 | 9,308 |
| K3 | Mazzaglia *et al.* (2012) | *A. deliciosa* | Korea, Jeonnam | 1997 | KN.2 | 962 | 10,103 |
| K4 | This paper | *A. chinensis* 'Hort16A' | Korea | 2014 |  | 258 | 46,118 |
| K5 | This paper | *A. chinensis* 'Hort16A' | Korea, Jeju | 2014 |  | 313 | 46,355 |
| K6 | This paper | *A. chinensis* 'Hort16A' | Korea | 2011 |  | 270 | 43,774 |
| K7 | This paper | *A. chinensis* 'Hort16A' | Korea | 2014 |  | 330 | 46,153 |
| K26 | McCann *et al.* (2013) | *A. chinensis* | Korea, Jeonnam | 1997 | KACC10584 | 290 | 36,930 |
| K27 | McCann *et al.* (2013) | *A. chinensis* | Korea, Jeonnam | 1998 | KACC10594 | 413 | 25,076 |
| K28 | McCann *et al.* (2013) | *A. chinensis* | Korea, Jeonnam | 1997 | KACC10574 | 297 | 37,347 |
| NZ13 | McCann *et al.* (2013) | *A. deliciosa* 'Hayward' | New Zealand, Te Puke | 2010 |  | 1 |  |
| NZ31<sup>1</sup> | Butler *et al.* (2013) | *A. deliciosa* | New Zealand, Paengaroa | 2010 |  | 382 | 33,149 |
| NZ32<sup>1</sup> | Butler *et al.* (2013) | *A. chinensis* | New Zealand, Te Puke | 2010 |  | 367 | 31,162 |
| NZ33<sup>1</sup> | Butler *et al.* (2013) |  | New Zealand, Te Puke | 2011 | TP1 | 380 | 31,549 |
| NZ34<sup>1</sup> | Butler *et al.* (2013) |  | New Zealand, Te Puke | 2011 | 6.1 | 386 | 46,155 |
| NZ35 | This paper | *Actinidia* sp. | New Zealand, Te Puke | 2010 |  | 330 | 47,026 |
| NZ37 | This paper | *Actinidia* sp. | New Zealand, Te Puke | 2010 | BF | 317 | 46,211 |
| NZ38 | This paper | *A. deliciosa* | New Zealand, Te Puke | 2014 | 627 | 322 | 46,155 |
| NZ39 | This paper | *A. deliciosa* | New Zealand, Te Puke | 2014 | 670 | 315 | 46,155 |
| NZ40 | This paper | *A. deliciosa* | New Zealand, Te Puke | 2014 | 793 | 322 | 45,201 |
| NZ41 | This paper | *A. deliciosa* | New Zealand, Te Puke | 2014 | 854.2 | 322 | 46,356 |
| NZ42 | This paper | *A. deliciosa* | New Zealand, Te Puke | 2014 | 632.1 | 322 | 48,669 |
| NZ43 | This paper | *A. deliciosa* | New Zealand, Te Puke | 2014 | 694.1 | 322 | 44,416 |
| NZ45 | This paper | *A. deliciosa* | New Zealand, Te Puke | 2014 | 1014 | 364 | 48,150 |
| NZ46 | This paper | *A. chinensis* | New Zealand, Matakana Island | 2012 |  | 325 | 48,671 |
| NZ47 | This paper |  | New Zealand, Te Puke | 2014 | 851 | 320 | 46,212 |
| NZ48 | This paper | *A. chinensis* 'Hort16A' | New Zealand, Te Puke | 2013 | 821 | 320 | 48,955 |
| NZ49 | This paper |  | New Zealand, Te Puke | 2011 | 691 | 325 | 46,155 |
| NZ54 | This paper | *A. chinensis* | New Zealand, Pukekohe | 2014 |  | 323 | 48,754 |
| NZ59 | This paper |  | New Zealand | 2015 |  | 315 | 44,129 |
| NZ60 | This paper |  | New Zealand | 2015 |  | 332 | 38,090 |
| P1 | Mazzaglia *et al.* (2012) | *A. deliciosa* 'Summer' | Portugal | 2010 | 346 | 354 | 31,352 |

<sup>1</sup>Simulated reads were generated from contigs available for these previously sequenced genomes
<sup>2</sup>Different sequencing runs were employed for draft genome assembly and variant calling

**Table S2.**
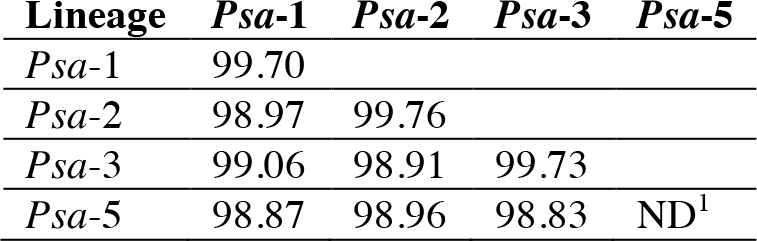
Average percent identity within and between Psa lineages

**Table S3.**
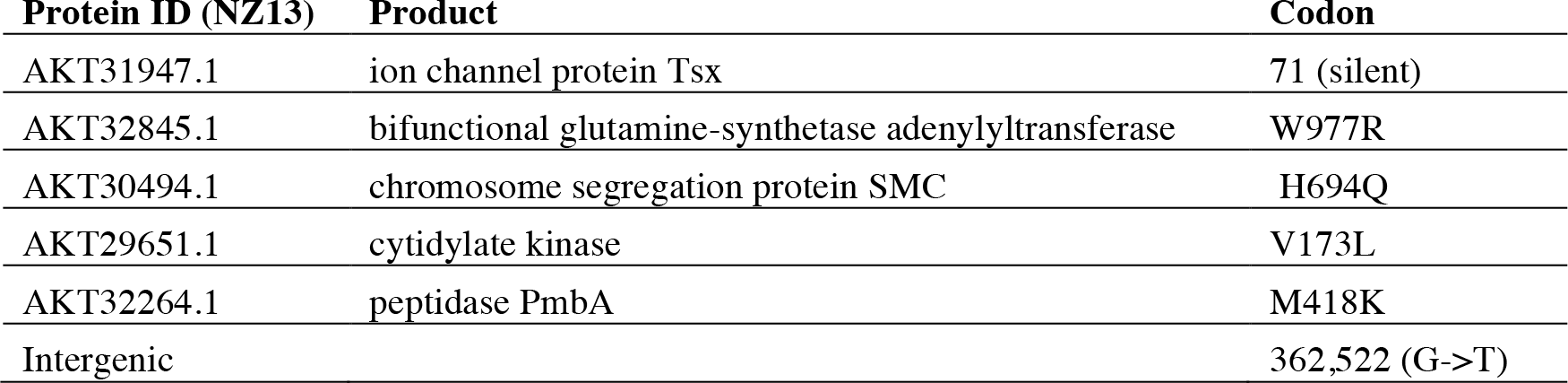
SNPs shared between all pandemic NZ and Japanese isolates

**Table S4.**
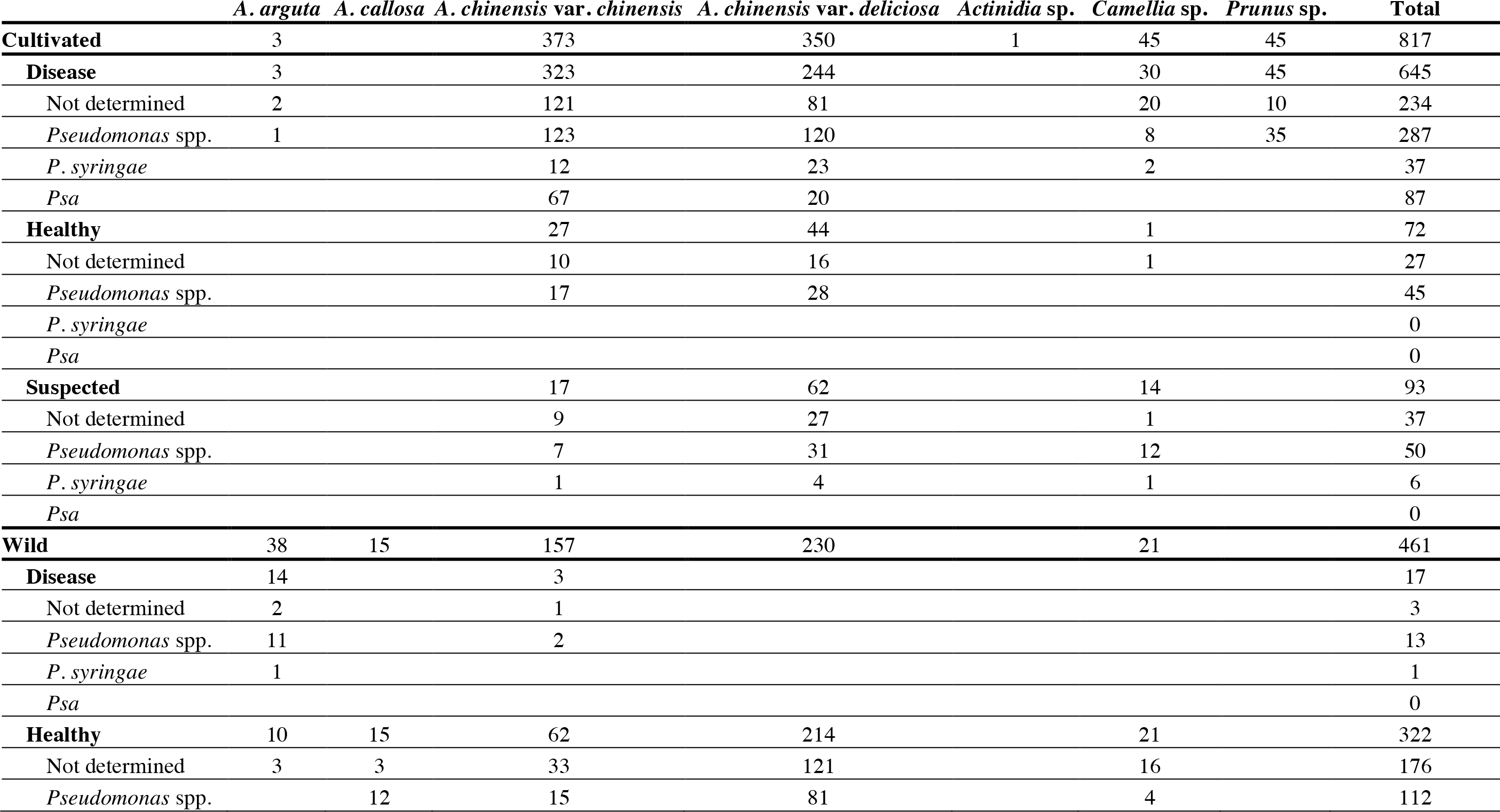
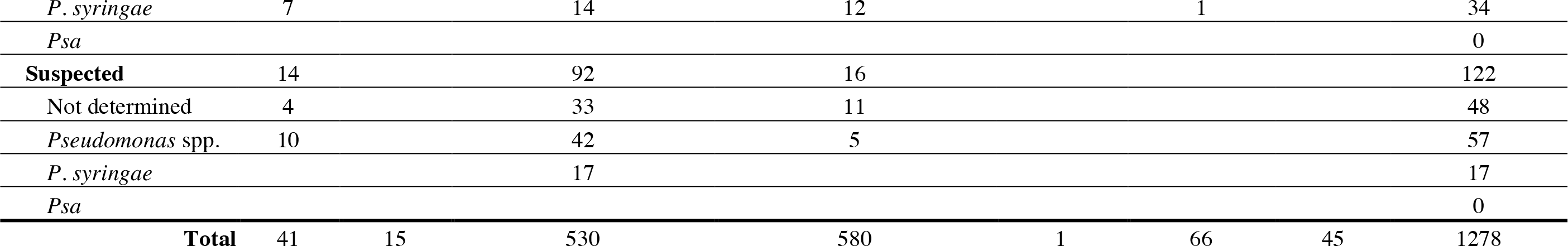
Isolates identified by cultivation and disease status of host

## FIGURES

**Figure S1.**
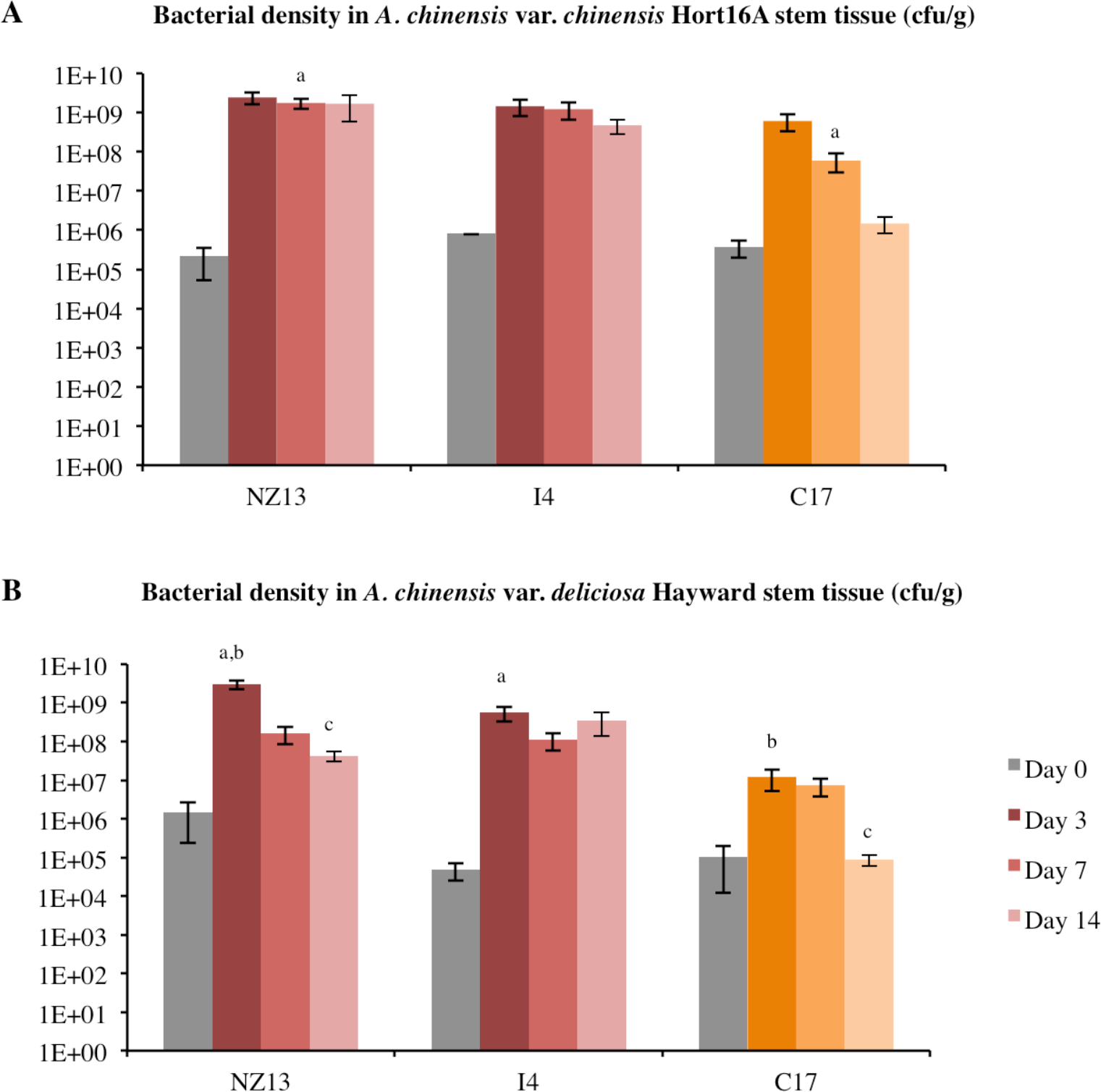
Bacterial growth assay on *A. deliciosa*. Bacterial growth of pandemic *Psa* NZ13, I4 (red) and divergent C17 (orange) strains on *A. chinensis* var. *chinensis* ‘Hort16A’ and *A. chinensis* var. *deliciosa* ‘Hayward’. Mean *in planta* bacterial density in stem tissue (cfu/g) at 0, 3 and 7 days post-inoculation is shown (mean ± SEM) with superscript denoting significant difference between strains at each sampling time (P<0.05, two-tailed t-test, unequal variance). Four replicate plants were assayed at day 0, and six replicates at each subsequent time point.

**Figure S2.**
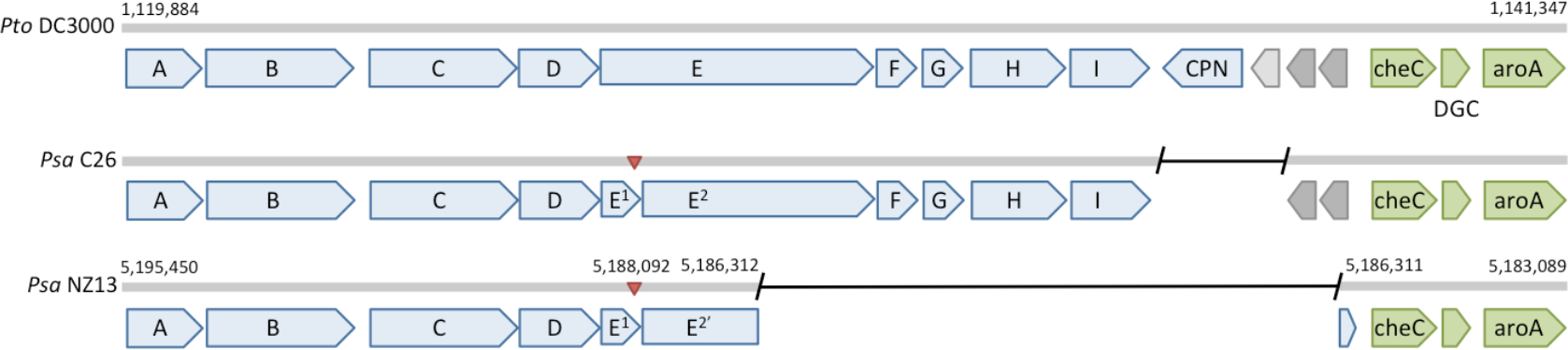
Wss operon disruption in *Psa*-3. Genes encoding components of the *wss* operon (blue), hypothetical and conserved hypothetical (light and dark grey), chemotaxis, diguanylate cyclase and *aroA* (green). Deletions (black line) and position of single base pair insertion (red triangle) displayed with reference to *Pto* DC3000. Insertion results in frameshift mutation in *wssE*, two predicted derivatives annotated as *wssE1* and *wssE2*. The subsequent 6.5kb deletion in the ancestor of the pandemic subclade results in the truncation of *wssE2*, annotated as *wssE2’*.

